# Ultrafast comparison of personal genomes

**DOI:** 10.1101/130807

**Authors:** Gustavo Glusman, Denise Mauldin, Leroy Hood, Max Robinson

**Author notes:** Correspondence, Gustavo Glusman Institute for Systems Biology, 401 Terry Ave N, Seattle, WA 98109, USA.

## Abstract

We present an ultra-fast method for comparing personal genomes. We transform the standard genome representation (lists of variants relative to a reference) into ‘genome fingerprints’ that can be readily compared across sequencing technologies and reference versions. Because of their reduced size, computation on the genome fingerprints is fast and requires little memory. This enables scaling up a variety of important genome analyses, including quantifying relatedness, recognizing duplicative sequenced genomes in a set, population reconstruction, and many others. The original genome representation cannot be reconstructed from its fingerprint; the method thus has significant implications for privacy-preserving genome analytics.

## Background

Personal genome sequences contain the information required for assessing genetic risks, matching genetic backgrounds between cases and controls in medical research, detecting duplicate individuals or close relatives for medical, legal, or historical reasons. Research purposes served by personal genome sequencing include classifying individuals by population, reconstructing human history, assessing and controlling the quality of the sequence information itself, computing kinship matrices to support genome-wide association studies (GWAS), and combining data sets for meta-analysis.

Many of these applications involve comparison of two or more personal genomes. However, the size, complexity, and diversity of representations in which they are stored makes comparison of personal genomes in their existing forms error-prone and slow, and therefore challenging to scale from pairs to the hundreds, thousands, or millions of individuals we will soon wish to compare in order to provide improved, personalized medical care.

Comparison of personal genomes requires cross-referencing on a very large number (potentially millions) of variants; this process can be slow enough to make the computation of all pairwise comparisons take a prohibitive amount of time. Current approaches to rapid genome comparison rely on severely limiting the number of single nucleotide polymorphisms (SNPs) to compare. A carefully curated set of 24 autosomal exome SNPs [1] captures up to 38 bits of information per genome and thus limits utility to establishing identity and is susceptible to failure when data are partially missing. A larger set of 1500 pre-selected SNPs has been used to match genomes directly from BAM files [2], with the requirement that these be mapped to the same reference genome. Similarly, focusing on tens of thousands of pre-selected SNVs enables multiple fast quality control computations in pedigrees [3]. Another powerful strategy involves deep compression of the data, coupled with advanced algorithms for fast querying and retrieval [4, 5]. The recent UNICORN method enables independent mapping of individuals onto ancestry spaces based on detailed maps of minor allele frequency distributions [6]. These strategies offer better speed at the expense of limited applicability, severely reduced accuracy, or strong reliance on detailed prior knowledge about the population being studied.

Perhaps more importantly, the diversity of representations of personal genomes remains a problem for each of these methods, which must translate SNPs represented in different ways to a common representation in order to assemble the data to compare. This diversity of representations arises from the variety of technologies used to determine personal genomes, their processing with diverse software pipelines and their representation relative to multiple versions of the reference genome. We explore each of these below.

The billions of base pairs in an individual’s genome can be assayed using a variety of techniques, the most complete of which is whole-genome sequencing (WGS). Several WGS technologies have been used to determine individual genomic sequences, differing in experimental protocols and in specific parameters such as read length and error tolerances; the resulting data are processed using diverse computational pipelines. Each technology brings a different set of observational biases and error modalities; the resulting observations are expressed in a diversity of ‘native’ vendor-specific file formats, and translated to one or more standard formats (e.g., variant call format, or VCF) in a variety of ways, leading to a bewildering diversity of flavors of semi-standard formats.

The resulting sequence is compactly represented as a list of differences (variants) from a reference genome. Due to the ongoing progress in improving the quality of the human reference genome, there are now several different versions (also known as ‘freezes’) in use. To date, many thousands of personal genomes have been ascertained, most of them expressed relative to the GRCh37 (also known as ‘hg19’) and GRCh38 (‘hg38’) freezes, though much legacy data exists using GRCh36 (‘hg18’) coordinates and even earlier versions. The position of a genomic variant depends on the reference version used: from one genome freeze to another, typically only the position of each variant along the chromosome changes, though in a small fraction of the cases a variant may be located in different chromosomes. Any two genomes of interest may be expressed relative to different reference genomes, necessitating conversion prior to comparison and analysis.

Even relative to the same reference version, a variant may have more than one representation, necessitating additional conversion steps potentially leading to errors. Each chromosome has multiple names (e.g., ‘NC_000003.12’, ‘chr3’ or just ‘3’ are all used refer to the third largest human chromosome), and different representations count the first nucleotide of each chromosome as position zero or one. Even with the same naming and numbering conventions, some variants can be expressed in more than one way, requiring normalization [7].

Sequencing the same genome using different technologies can yield differing results, as each technology has its own biases. Even when using the same technology, reference and encoding, sequencing the same genome repeatedly can give somewhat different results due to the stochastic nature of genome sequencing, to batch effects, or to differences in the computational pipelines used.

Looking to the future, additional (long-read) technologies will enable *de novo* genome assembly to become commonplace; the reference genome representation will change to a graph format, further breaking the concept of absolute coordinates; and the number of genomes available will soon be in the millions. These trends will deepen, not lessen the complexity of genome comparison.

We present here a new, rapid method for summarizing personal genomes that does not require knowledge of the technology, reference and encoding used, and yields ‘genome fingerprints’ that can be used to facilitate many standard problems in genomics. Thanks to their reduced size, computation on the genome fingerprints is orders of magnitude faster and requires little memory, enabling comparison of much larger sets of genomes. No individual variants or other detailed features of the personal genome can be reconstructed from the fingerprint, thereby allowing private information to be more closely guarded and protected by decoupling genome *comparison* from genome *interpretation*. Fingerprints of different sizes allow balancing the speed and accuracy of the comparisons, and due to the high value of estimating relatedness, the potential applications of genome fingerprinting range from basic science (study design, population studies) to personalized medicine, forensics and genealogy research.

## Methods

### Overview

Our algorithm summarizes a personal genome as a ‘fingerprint’ (Figure 1). Conceptually, any variant-oriented representation of a personal genome (e.g. a VCF file or vendor-specific format, regardless of encoding or reference genome used) includes a list of variants, including their position and reference and alternative alleles, sorted by position. A ‘raw’ fingerprint is a tally of consecutive biallelic single-nucleotide variants (SNV) pairs grouped on a combination of these two attributes. We then normalize the raw fingerprint to account for systematic differences in frequency between groups by allele and by position. The resulting ‘normalized’ fingerprint preserves differences at the species level, e.g. between individuals from different populations. Averaging the normalized fingerprints of the individuals in a population yields a ‘population’ fingerprint, which can be subtracted from an individual’s normalized fingerprint to produce a ‘population-adjusted’ fingerprint suitable for more sensitive detection of related genomes.

**Figure 1.**
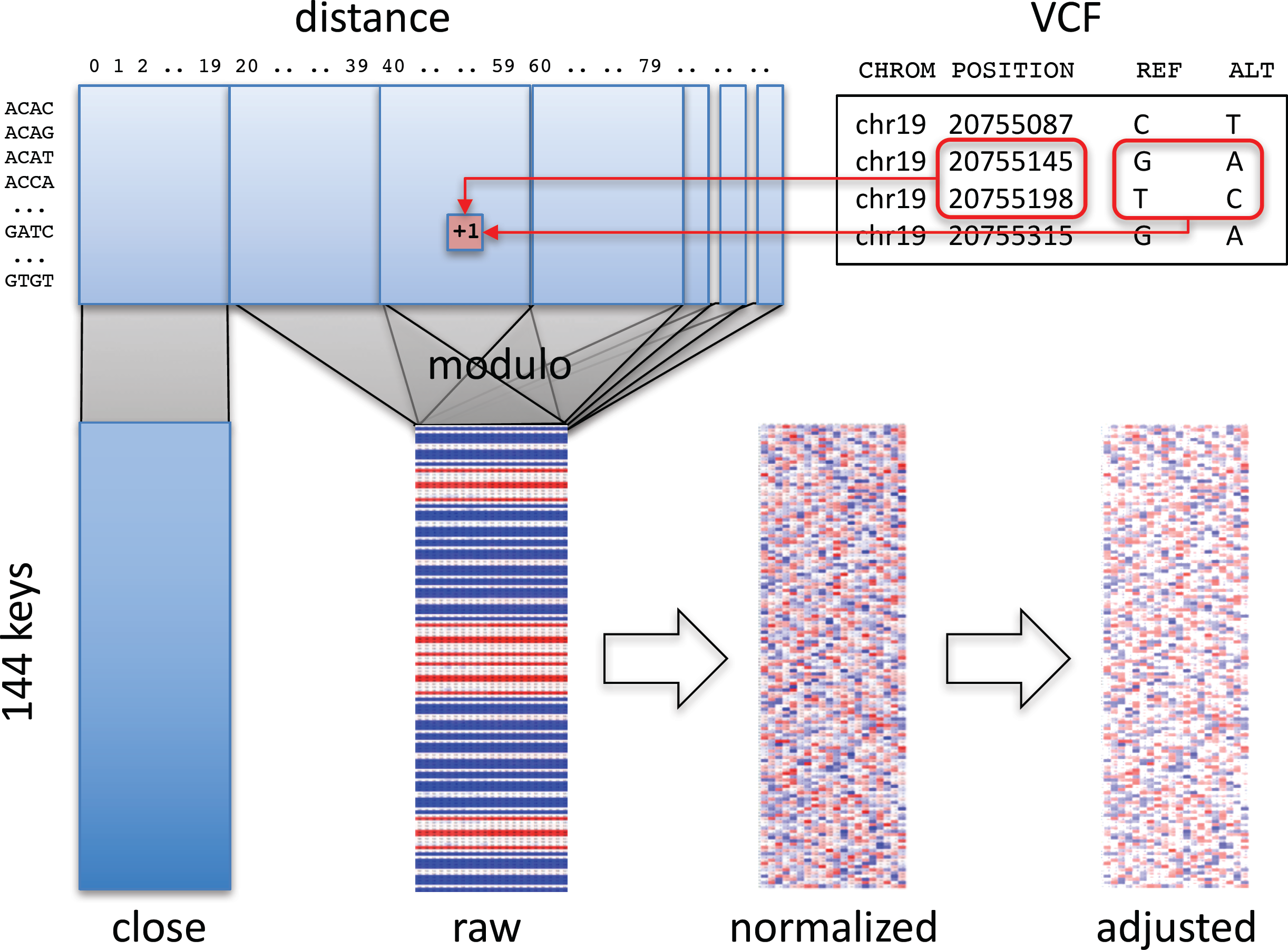
Overview of method. Pairs of consecutive SNVs in the input file (upper right) are encoded into a table (upper left) by SNV key and by distance. A section of the table, informative about technology, is segregated (‘close’ matrix, lower left). The rest of the table is folded using the modulo function to generate the raw fingerprint (‘raw’ matrix, lower center), which is then normalized and adjusted to the closest population (lower right).

### Computation of raw fingerprints

The first stage in computing genome fingerprints yields a ‘raw’ fingerprint, a 144 x *L* table of SNV pair counts (*L*, the fingerprint length, is the main parameter of the method, defaulting to 20). We classify each pair of consecutive SNVs by the combination of reference and alternate alleles at both SNVs: this information determines the row in the table. We also consider the distance between the SNVs: this information determines the column in the table. When studying genomes produced using Complete Genomics and Illumina technologies, we observed differences in the distribution of distances between consecutive SNVs, for short distances (Supp. Figure 1). These differences largely reflect each vendor pipeline’s encoding of multi-nucleotide variants; some pipelines represent these as individual SNVs, potentially leading to interpretation errors [8]. We thus exclude SNV pairs that are separated by fewer than *C* base pairs (defaulting to 20) from the final fingerprint. To avoid technology-specific artifacts, we recommend *C* at least 5 (see below).

To compute a raw fingerprint:

1) Parse the autosomes in the personal genome (e.g., from a VCF file) to identify biallelic SNVs as variants with reference allele of length 1 and single alternate allele of length 1, both in the case-insensitive alphabet [A C G T]. Ignore all other types of variants: two biallelic SNVs separated by other types of variants are still considered consecutive.

2) For each SNV, join its reference and alternative alleles into one of 12 possible SNV keys. For example, reference allele ‘G’ and alternative allele ‘A’ form key ‘GA’.

3) Join the keys of consecutive SNVs on the same chromosome to form one of 144 possible SNV pair keys; for example, consecutive SNVs with keys ‘GA’ and ‘TC’ form the SNV pair key ‘GATC’.

4) Compute the number of base pairs between the two SNVs (the ‘distance’ between the SNVs along the reference genome). If this distance is smaller than *C*, increment by one the corresponding value in the [144 x *C*] ‘close’ matrix (Figure 1), and skip the next steps.

5) Compute a reduced distance as the distance between the two SNVs, modulo the *L*.

6) Increment by one the corresponding value in the [144 x *L*] ‘raw’ matrix.

### Normalization of fingerprints

The second stage in computing genome fingerprints performs a normalization of the raw fingerprint to account for the different frequencies of transitions and transversions, distance and parameter effects (Supp. Figure 2). The normalized fingerprints do not show any remaining structure (Supp. Figure 3).

The normalization is performed in two steps:

1) Normalization by distance: subtract the mean and divide by the standard deviation of each column.

2) Normalization by SNV pair key: subtract the mean and divide by the standard deviation of each row.

### Population fingerprints

We compute a *population fingerprint* as the average of the normalized fingerprints from the individuals in the population. These fingerprints must have been computed using the same parameters (*C* and *L*).

### Adjustment to population

We compute a *population-adjusted fingerprint* for an individual by subtracting a population fingerprint from the normalized fingerprint of that individual. The individual fingerprint and the population fingerprint must have been computed using the same parameters (*C* and *L*). Similar to normalized fingerprints, population-adjusted fingerprints show no internal structure (Supp. Figure 4).

### Fingerprint comparison

To compare two fingerprints, concatenate the rows of each fingerprint matrix into a vector and compute the Spearman correlation between the two vectors. This same procedure is appropriate for comparing two normalized fingerprints or two population adjusted fingerprints, whether adjusted to the same or different populations.

### Binary fingerprints

We also evaluated a minimal version of the genome fingerprints, in the shape of a binary string of length 144, as follows (Supp. Figure 5). We (a) compute a raw fingerprint matrix with *L*=2, and (b) for each row in the matrix, corresponding to a SNV pair key, add to the binary fingerprint a 1 if the value of the second column is larger than the value in the first column. There is no need to normalize binary fingerprints. To compare two binary fingerprints, we count how many bits are identical between the two, divide by the number of bits (144) and square the resulting fraction.

### Evaluation of resilience of genome fingerprints

#### Different reference versions

To evaluate the resilience of the fingerprinting method to different versions of the reference genome, we computed fingerprints (*L*=20 and binary) for a set of 69 genomes sequenced by Complete Genomics, Inc. (http://www.completegenomics.com/public-data/69-genomes/); these genomes were mapped to both GRCh36 (hg18) and GRCh37 (hg19). We then computed all pairwise correlations, which include a) each genome on both references, b) combinations of different genomes on the same reference and c) combinations of different genomes with different references.

#### Format and normalization

To evaluate the resilience of the fingerprinting method to transformation from vendor-specific formats to standard representations, we studied 2436 CGI genomes delivered in ‘var’ format and 1618 CGI genomes delivered in ‘masterVar’ format. The var format reports each allelic observation in a separate line, while the masterVar format represents each locus in one line (in similarity with the VCF format). We used custom parsers to transform these vendor-specific genome representations to VCF format, and normalized them using vt normalize [7]. We then compared genome fingerprints computed from the original (var or masterVar) representations and from the normalized VCF representations. We performed these computations with *L*=20 and with binary fingerprints.

#### Significant filtering and post-processing

To evaluate the resilience of the fingerprinting method to complex post-processing procedures, we studied 154 genomes from the 1000 Genomes Project for which individual WGS data are available from the International Genome Sample Resource (http://www.internationalgenome.org/announcements/complete-genomics-data-release-2013-07-26/). The genomes were analyzed using Complete Genomics’ analysis pipeline versions 2.2.0.19 through 2.2.0.26 and reported in masterVar format. We compared for each genome fingerprints computed from the masterVar representation and from the version extracted from the multi-sample VCFs in release 20130502 (filenames: ALL.chrNN.phase3_shapeit2_mvncall_integrated_v5.20130502.genotypes.vcf.gz). We performed these computations with *L*=20, *L*=200, and binary fingerprints.

#### Sequencing technology effects

To evaluate the resilience of the fingerprinting method to differences between technologies, we computed fingerprints for several versions of the “platinum” NA12878 genome, some of them sourced from the Genome In A Bottle consortium (downloaded from ftp://ftp-trace.ncbi.nlm.nih.gov/giab/ftp/data/NA12878/analysis/). These assemblies were sequenced using multiple versions of CGI’s technology and Illumina, processed using various pipeline versions, mapped to GRCh36, GRCh37 and GRCh38 (Figure 2). We performed these comparisons with *L*=5, *L*=20, *L*=120, and binary fingerprints.

**Figure 2.**
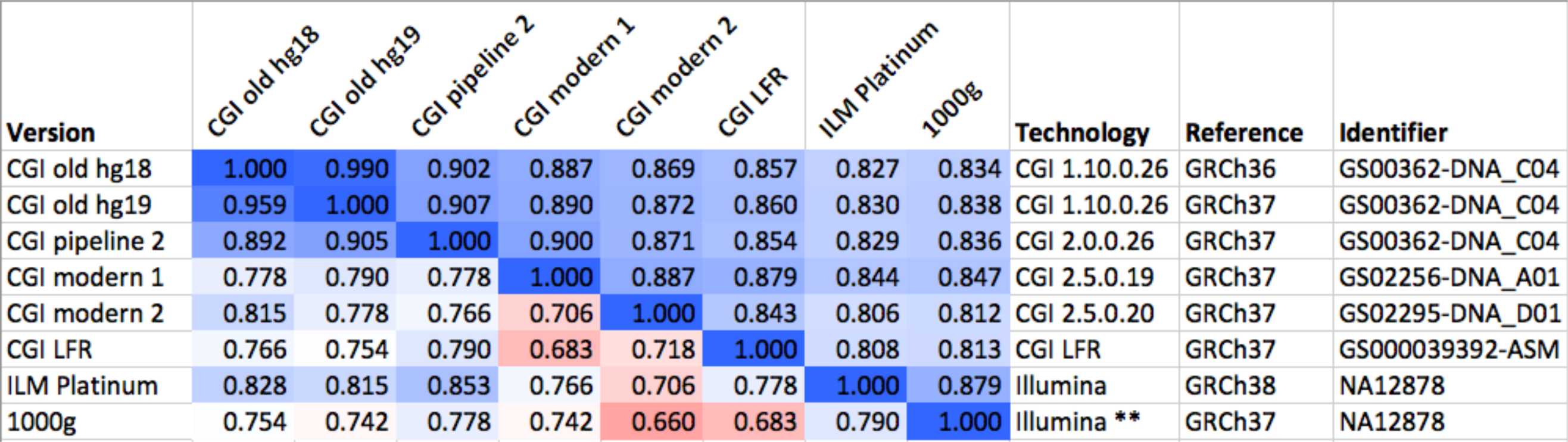
Comparison of several versions of the same genome. We compared six versions of the genome of the same individual (NA12878), one of them (GS00363-DNA_C04) processed in three different ways. The values above the diagonal of the matrix represent pairwise Spearman correlations between fingerprints with *L*=20, while numbers under the diagonal represent comparisons of binary fingerprints. Color scale from 0.5 (red) through 0.75 (white) to 1.0 (blue). CGI: Complete Genomics, Inc. ILM: Illumina. LFR: long fragment read. 1000g: 1000 Genomes Project. ** highlights the post-processing of the Illumina-based genome sequence as done by the 1000 Genomes Project.

#### Missing data

To evaluate the resilience of the fingerprinting method to missing data, we performed a simulation in which we degraded a genome to increasing degrees and compared the resulting fingerprints to the original. We computed a series of fingerprints for the same genome by varying the probability of not observing each individual variant, excluding 1, 5, 10, 15, 20, 25, 30, 35, 40, 45, 50, 60, 70, 80, 90, 95, and 99% of the variants at random (Figure 3). We performed this simulation with *L*=10, *L*=20, *L*=50, *L*=120, and binary fingerprints.

**Figure 3.**
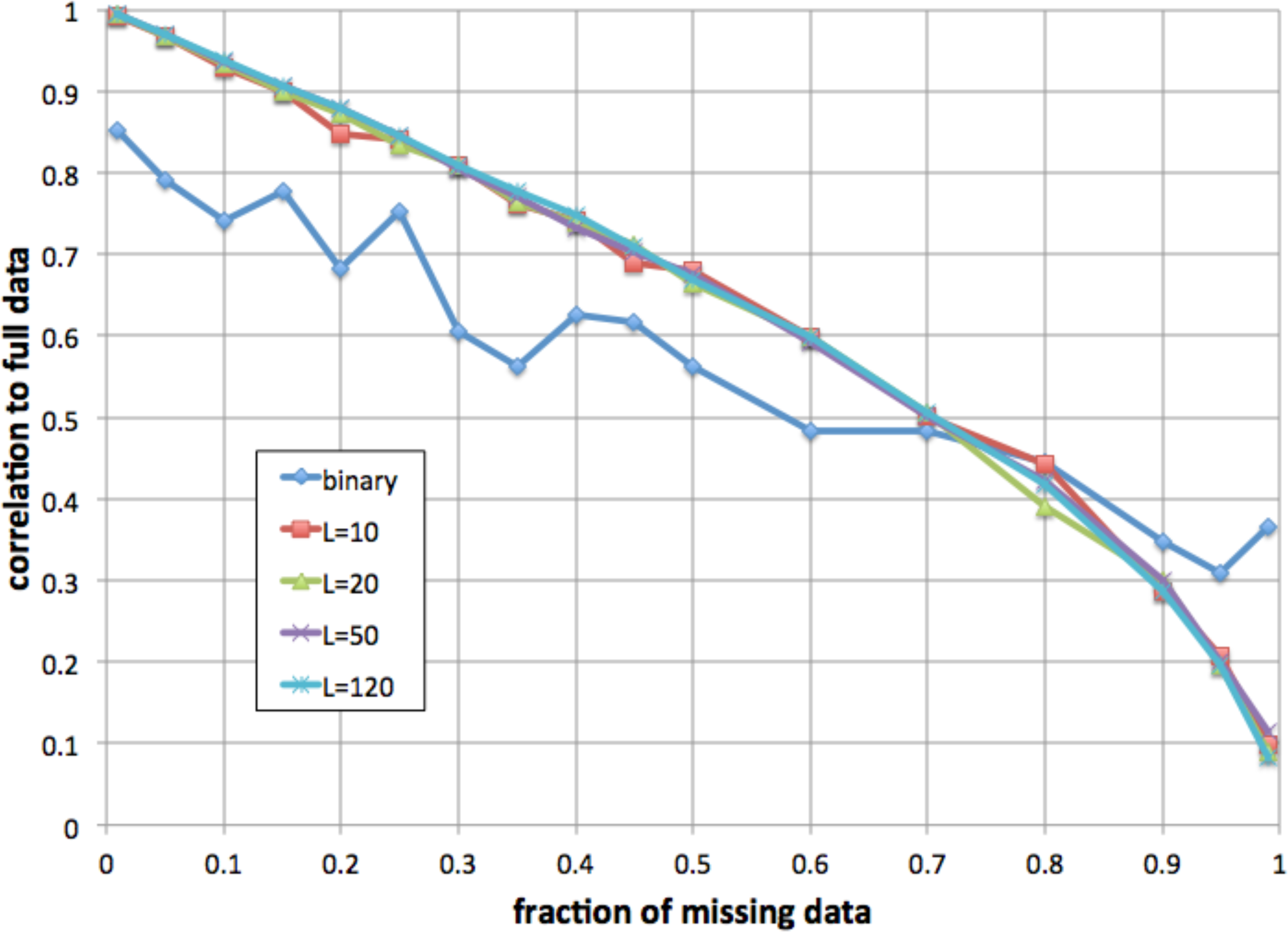
Resilience to missing data. We simulated increasing fractions of missing data and computed the correlation between the resulting fingerprint and the fingerprint derived from the full data. For fingerprints with *L*=10, *L*=20, *L*=50, and *L*=120, the method was resilient to up to 30% data loss. The binary fingerprint was resilient to up to 5% data loss.

### Pedigree study

We studied WGS data from 35 individuals in a large family (Figure 4). All genomes were sequenced by Complete Genomics, Inc. from blood (n=25) or saliva (n=10) samples, and processed using pipeline version 2.5.0.20. The genome data and a description of the family pedigree were donated by the private family. We categorized all pairwise relationships within the family as sibling, parent/child, half-sibling, aunt/uncle, grandparent, cousin, second cousin, second cousin once removed, unrelated, and other. The last category included more complex relations, e.g., child and grandchild of half-siblings. We computed for each individual a series of genome fingerprints using *L* in the range 2-200. For each *L* we computed all pairwise correlations, then computed their average and standard deviation stratified by the relationship categories listed above.

**Figure 4.**
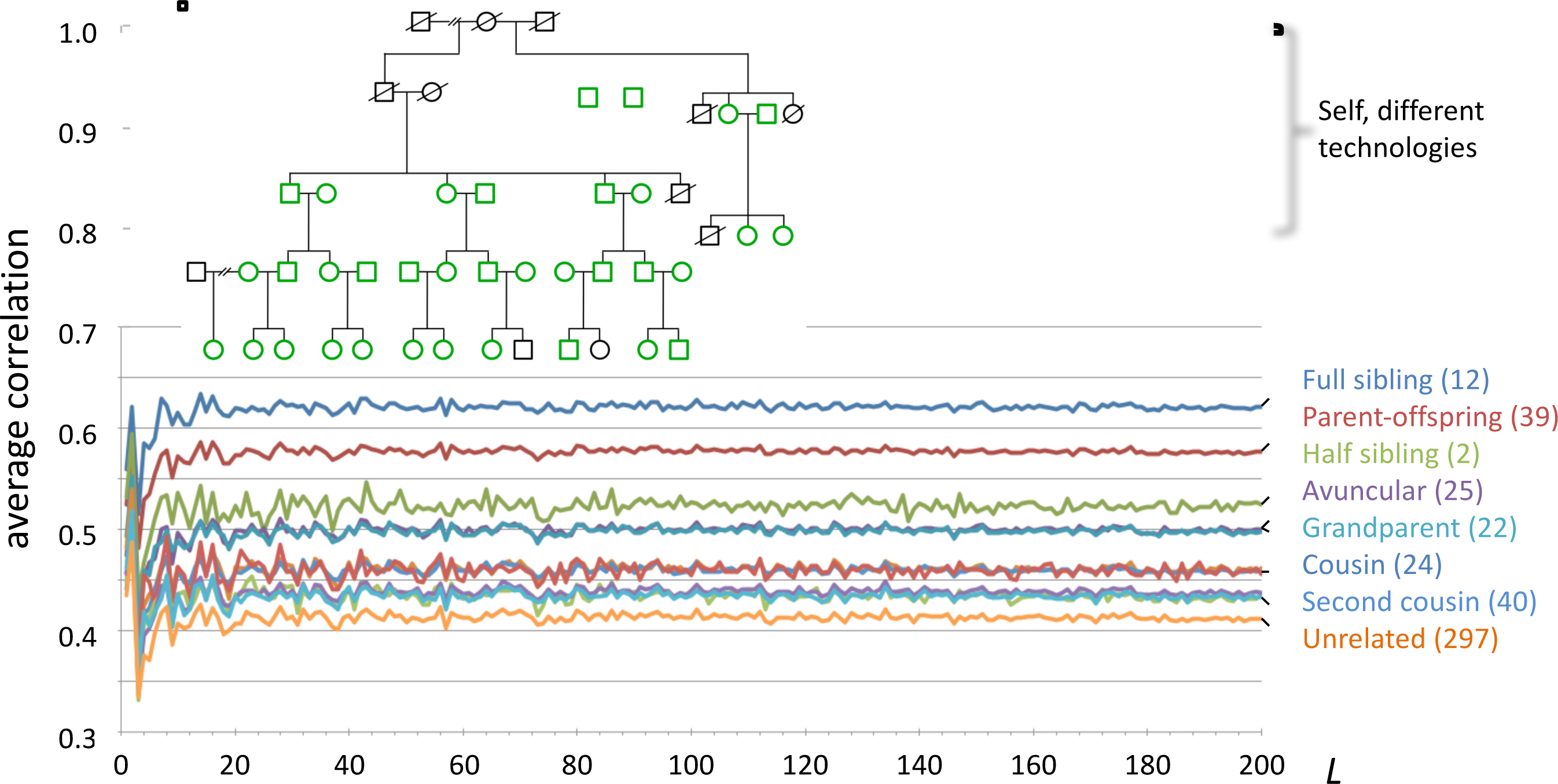
Fingerprint correlation reflects degree of relatedness. Each trace represents the average correlation between fingerprints of individuals in each relatedness group, as a function of *L*. The number of pairs in each class is shown between parentheses. Inset: family structure; green icons represent individuals whose genomes were sequenced. Not shown: 169 additional pairs with complex relationships.

### Population reconstruction

We computed fingerprints (*L*=20, *L*=120, and binary) for each of the 2504 genomes in the 1000 Genomes Project data set from the version extracted from the multi-sample VCFs in release 20130502 (filenames:

ALL.chrNN.phase3_shapeit2_mvncall_integrated_v5.20130502.genotypes.vcf.gz). For comparison, we analyzed the same data by principal components analysis (PCA) as follows.

We identified SNPs with a minor allele frequency of 5% or more, removed SNPs in complete linkage disequilibrium with a SNP to the left (i.e. a smaller chromosomal position), retained 5% at random (298,454 SNPs) and counted occurrences of the minor allele (0, 1, or 2) in each genome to form a 2504 x 298,454 genotype matrix M. We performed PCA using the R function call prcomp(M,center=TRUE,scale=TRUE). (Figure 5)

**Figure 5.**
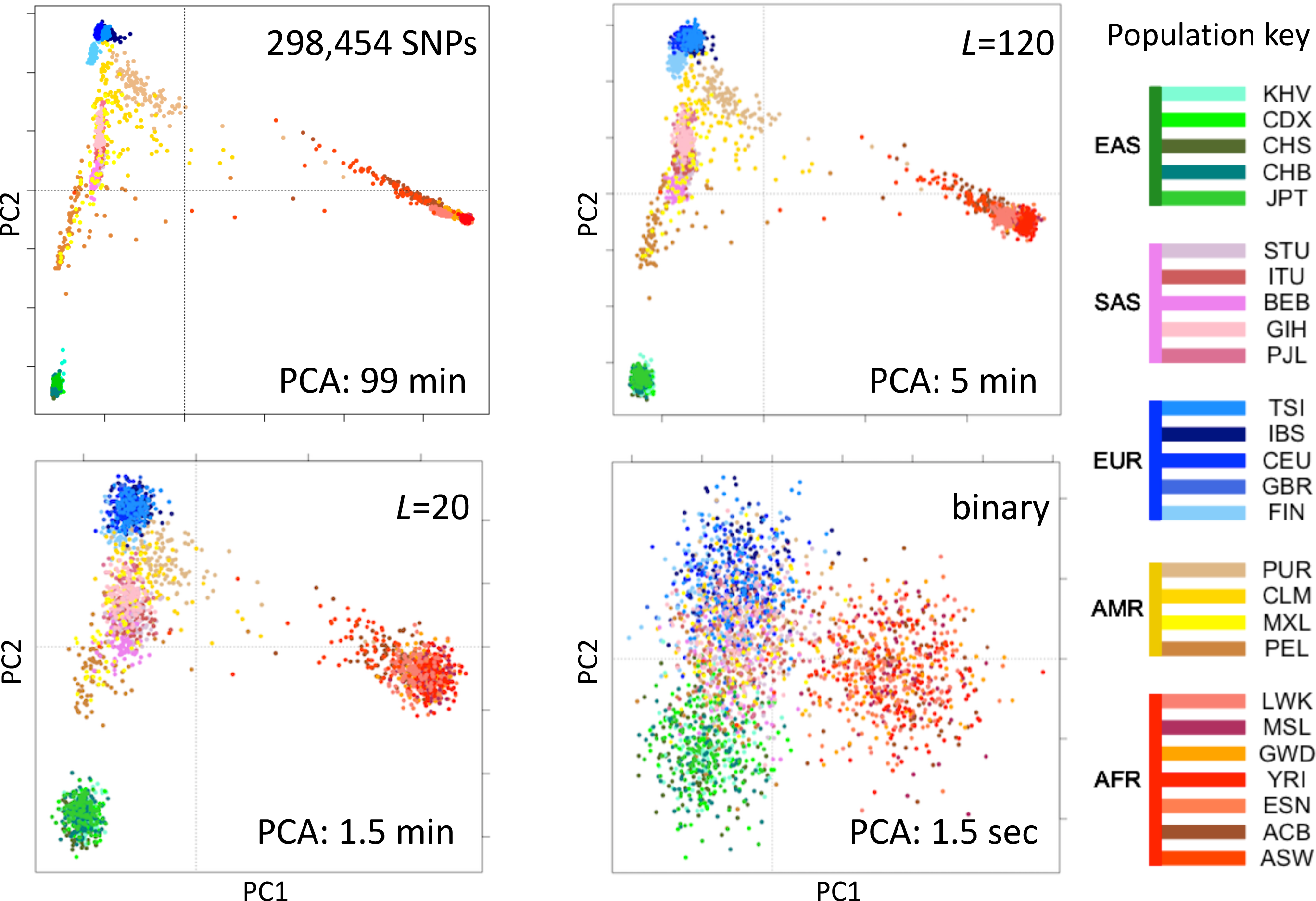
Reconstruction of population structure of the 1000 Genomes Project data set. Each panel shows the principal component analysis (PCA) of the 2504 individuals using ∼300,000 SNPs (upper left), genome fingerprints with *L*=120 (upper right), *L*=20 (lower left), and binary fingerprints (lower right). Individuals are color coded according to their population as per the key to the right. EAS, SAS, EUR, AMR and AFR: East Asian, South Asian, European, Admixed American and African, respectively. PC1 and PC2: first two principal components.

### Evaluation of fingerprints for population classification

We computed a population fingerprint for each of the annotated populations studied by the 1000 Genomes Project. To identify the population closest to each individual, we compared each individual’s normalized fingerprint to the population fingerprints (using Spearman correlation, as described for comparing individual fingerprints). Each individual was considered classified as belonging to the closest population. To avoid bias, we excluded each individual from the computation of their own population fingerprint.

### Population-adjusted fingerprints

We computed a population-adjusted fingerprint for each individual in the 1000 Genomes data set by subtracting the population fingerprint (excluding the individual) from the individual’s normalized fingerprint. We computed all pairwise Spearman correlations and identified outliers using the absolute deviation around the median [9] with false discovery rate of 5%.

## Results

### A novel genome encoding method

We developed a novel algorithm for computing ‘fingerprints’ from genome data; the algorithm is akin to locality-sensitive hashing [10]. These fingerprints are rapidly computed, only need to be computed once per genome (not once per comparison), and can be rapidly compared to determine whether two genome sequences are derived from the same individual, closely related individuals, or unrelated individuals. Moreover, the original genome representation cannot be reconstructed from the fingerprint, and fingerprints can be shared when privacy concerns prevent sharing the genome itself.

Importantly, the fingerprints can be computed starting from any of several file formats, with a variety of encodings and relative to any reference version, and the resulting fingerprints are directly comparable without further conversion.

Based on characteristics of consecutive pairs of SNVs (see Methods), our algorithm computes two small matrices of numbers for each genome (Figure 1), one of which (‘close’) holds potentially useful information about the technology used to generate the genome, and the other (‘raw’) represents the genome itself. The latter is then normalized and potentially adjusted to the relevant population fingerprint. Each of these matrix versions has its own utility.

The main parameter of our algorithm, *L*, determines the size of the fingerprint. Smaller fingerprints (e.g., *L*=20) are useful for fast genome comparisons to determine identity, while larger fingerprints (e.g., *L*=120, *L*=200) retain more information and better support detailed analyses like population reconstruction.

### A minimal fingerprint

We further sought to create a minimal genome fingerprint. Using the smallest meaningful table (*L*=2), we created a fingerprint of just 144 bits for each genome. Each bit represents, for each of the 144 possible SNV pair keys (e.g., ‘GAGT’), whether the distance (the number of intervening base pairs) between the consecutive SNVs is more frequently an odd or an even number. This simple hash function yields a ‘binary barcode’ of a genome (Supp. Figure 5) with negligible storage and memory footprints, trivially comparable to similarly computed ‘barcodes’ by counting binary matches.

### Computation on genome fingerprints is fast

Computation of genome fingerprints is very fast, typically requiring 15-45 seconds per genome, depending on the file format. The computation requires a single-pass read of the genome and depends principally on the time it takes to read the file (I/O bound). It is also trivially parallelizable: computation of one genome fingerprint does not depend on the results of similar computations for other genomes. It is also possible to compute partial raw fingerprints from sections of the genome, e.g., individual chromosomes; such partial fingerprints can then be trivially summed yielding the raw fingerprint for the whole genome.

Thanks to the small size of the fingerprint matrices, fingerprint comparisons are extremely fast. We performed all-against-all comparisons in a set of 11,726 genomes. The nearly 69 million comparisons required 10 CPU hours (0.5 milliseconds per comparison) with *L*=20, and under half a CPU hour (25 microseconds per comparison) for binary fingerprints. Fingerprint comparisons are also independent and trivially parallelizable.

### Genome fingerprints are resilient

#### Fingerprints are resilient to reference versions

Mapping raw sequence data to different reference versions can yield disparate variant calls. The absolute coordinates for most observed variants will differ between references; a fraction of the variants will differ also in other aspects including reference allele, variant allele and even chromosome assignment and strand. We studied a set of 69 genomes that were mapped to two reference versions and evaluated the effect of reference version on fingerprint comparison. With *L*=20, the Spearman correlation between fingerprints derived from genomes mapped to GRCh36 and GRCh37 ranged from 0.989 to 0.991, while fingerprints derived from different genomes in this set ranged from 0.222 to 0.685 (see example in Figure 2). Binary fingerprints yielded lower correlations that nevertheless clearly separated self-comparisons (0.853-0.986) from comparisons of different individuals (0.243-0.742). Genomes mapped to different references therefore yield highly comparable fingerprints.

#### Fingerprints are resilient to format differences

Genome sequencing vendors may deliver results in non-standard file formats that best convey technology-specific characteristics of the data; the variants themselves may be represented in inconsistent ways. Considering two vendor-specific file formats (Complete Genomics’ var and masterVar), we evaluated the effect on fingerprinting of standardizing the genome data representation by transforming to the standard VCF format and by normalization of variants [7]. With *L*=20, the correlation between fingerprints derived from var files and from their normalized VCF counterparts was 0.9911 +/- 0.0008. For masterVar files, the correlation was 0.9982 +/- 0.0015. Binary fingerprints yielded marginally lower correlations: 0.9262 +/- 0.0313 for var files and 0.9688 +/- 0.0211 for masterVar files. We found that this transformation yields nearly identical fingerprints.

#### Fingerprints are resilient to complex post-processing

The 1000 Genomes Project applied a number of post-processing steps to the collection of 2504 genomes to normalize variant calls and exclude spurious or unreliable variants. Thus, for each genome in the data set there are at least two versions: the initial, unprocessed individual genome, and the harmonized version in the context of the large study. We evaluated whether fingerprint comparison is resilient to this complex post-processing procedure (see example in Figure 2). With *L*=20, the correlation between initial and post-processed versions was 0.798 +/- 0.010. We observed nearly identical results with *L*=200: 0.798 +/- 0.008. The lowest observed correlation for an individual genome was 0.776, well above the correlations observed for different individuals, even those closely related. As expected, binary fingerprints were more strongly affected by the post-processing procedure, yielding correlations of 0.660 +/- 0.057. Here, the lowest observed correlation for an individual was 0.463, significantly below the highest correlation observed between binary fingerprints of two related individuals (0.742).

#### Fingerprints are resilient to technology differences

We compared fingerprints computed from various versions of the same genome (NA12878) as ascertained using different technologies, mapped with different pipelines and using different references (Figure 2). Using fingerprints with *L*>=20, all comparisons yielded correlations higher than 0.8. Shorter fingerprints (*L*=5) yielded correlations above 0.75 (not shown). As expected, binary fingerprints were more sensitive to technology differences.

#### Fingerprints are resilient to missing data

We evaluated whether fingerprint comparison is resilient to missing data by simulation. For a variety of fingerprint sizes, we degraded a personal genome by excluding a fraction of the variants prior to computing fingerprints. For most fingerprint sizes, we observed a monotonic and equivalent decrease in Spearman correlation between the degraded genome fingerprints and the original, undegraded genome fingerprint. Given the 0.8 lowest correlation observed for different versions of the same genome, we conclude that the fingerprinting method is resilient to up to 30% missing data (Figure 3). As expected, the less informative binary fingerprint is less resilient (up to 5% missing data).

### Fingerprint similarity reflects degree of relationship

We compared pairwise genome fingerprint correlations with known family relationships in a large pedigree (Figure 4). The family relationships spanned the range from close (e.g., siblings) to distant (e.g., second cousins once removed) and also included unrelated individuals (joining the family by marriage). We observed that fingerprint correlations decrease with increasing degree of relationship. This demonstrates that our genome fingerprinting method is locality-sensitive. We also observed that *L*>=20 is sufficient for distinguishing most relationship levels on average. Larger fingerprints have lower standard deviations (Supp. Figure 6), improving resolution and the confidence of close relationship. The genomes of all individuals in this pedigree were sequenced using the same technology and processed using similar pipeline versions and the same reference version; this setup is typical for research studies. Combining data obtained using multiple technologies and processing pipelines could reduce the ability to confidently distinguish between relationship levels.

Of note, fingerprints derived from siblings tend to be more highly correlated than fingerprints of parents and their offspring. In both cases the degree of relationship is the same, but the identity by descent (IBD) pattern differs: parents and offspring share 100% of the genome in IBD1 state, while siblings share (on average) 50% of the genome in IBD1 state and 25% in IBD2 state. This increases the probability that sibling pairs will share heterozygous variants, especially heterozygous rare variants, as compared with parent/child pairs. For similar reasons, fingerprints of half siblings are more correlated than fingerprints of relatives of the same average degree (grandparental and avuncular relationships).

### Fast reconstruction of population structure

We tested the utility of genome fingerprints for population studies. We computed fingerprints for the 2504 individuals from 1000 Genomes Project and used PCA to reconstruct the known population structure (Figure 5) in a fraction of the time required to perform the same task using standard methods and with much smaller memory requirements. The quality of the reconstruction depended on fingerprint size: while binary fingerprints yielded only a first approximation, fingerprints with *L*=20 yielded good separation at the continental level, and fingerprints with *L*=120 yielded excellent population structure reconstruction. PCA applied to fingerprints of different values of *L* provided results highly correlated with results from PCA applied to variants, with convergence to the same principal component axes as either the number of variants or the *L* increased. For a sufficient amount of data in either form, correlation between corresponding principal components was > 0.99 for the first 5 to the first 10 components. This strongly suggests PCA of either kind of data provides equivalent information regarding population structure.

Importantly, the population reconstruction workflow using fingerprints is simple and requires no prior knowledge. The standard workflow requires creating multi-sample VCFs (which in turn requires normalizing variants), filtering out unreliable variants (which requires prior knowledge), filtering by allele frequency (requiring population frequencies), paying attention to linkage disequilibrium, making assumptions about variants not observed in some samples and, finally, selecting a suitably sized subset of variants to balance resolution with memory and computational requirements. In contrast, using genome fingerprints it is possible to reconstruct population structure by computing fingerprints directly from the unprocessed individual genomes and combining the fingerprints into a matrix ready for analysis with PCA or any other method of choice.

### Utility for fast and simple population assignment

Individuals from the same population share some evolutionary history, and therefore some of the SNV pairs counted in computing genome fingerprints. It is thus useful to summarize the fingerprints of a population, both to estimate the “center” of the population’s fingerprints and their variability around that center (population diversity). Such “population fingerprints” have a variety of uses, including population assignment for individuals. We implemented a simple and very rapid method in which we first compute fingerprints (*L*=120) for each population in the 1000 Genomes data set by simply averaging the fingerprints of the genomes in each population. We then compute the correlation between the fingerprint of a query genome and the fingerprint of each population: the genome is assigned to the population with which it is most strongly correlated. We tested this method by “leave one out” cross-validation. The correct population was identified as the best match for 2047 of 2504 query genomes (82% of cases). Some of the populations in the 1000 Genomes data set are very closely related and difficult to separate; for many practical purposes (e.g., selecting appropriate allele frequencies) using such closely related populations yields equivalent results. Accepting the 2nd or 3rd best population matches increased the success rate to 96% and 98%, respectively. At the continental resolution (5 regions: AFR, AMR, EAS, EUR, SAS), the best match was correct for all but 42 admixed AMR genomes.

### Utility of population-adjusted fingerprints

Fingerprints can be manipulated in a familiar way to perform different distance-based analyses. Population fingerprints, for example, are simple averages of a group that capture the features common to the group with respect to distance, but without corresponding interpretations that could be used for racial profiling or stereotyping. Population adjusted fingerprints allow analysis of relationships within a population, beyond the features shared among the population. This allows close relationships to be distinguished from membership in the same population and may help increase resolution of population structure reconstruction. We adjusted the *L*=120 fingerprints for the 2504 individuals in the 1000 Genomes data set relative to their stated population of origin and performed all pairwise comparisons (Figure 6). Of note, these correlation values among population adjusted fingerprints are not comparable with those observed among normalized fingerprints in the family study (Figure 4). The similarity between unrelated individuals derived from the same population (e.g., Figure 4) is removed by adjustment to the population average. Thus, population-adjusted fingerprints for unrelated individuals show no significant correlation.

**Figure 6.**
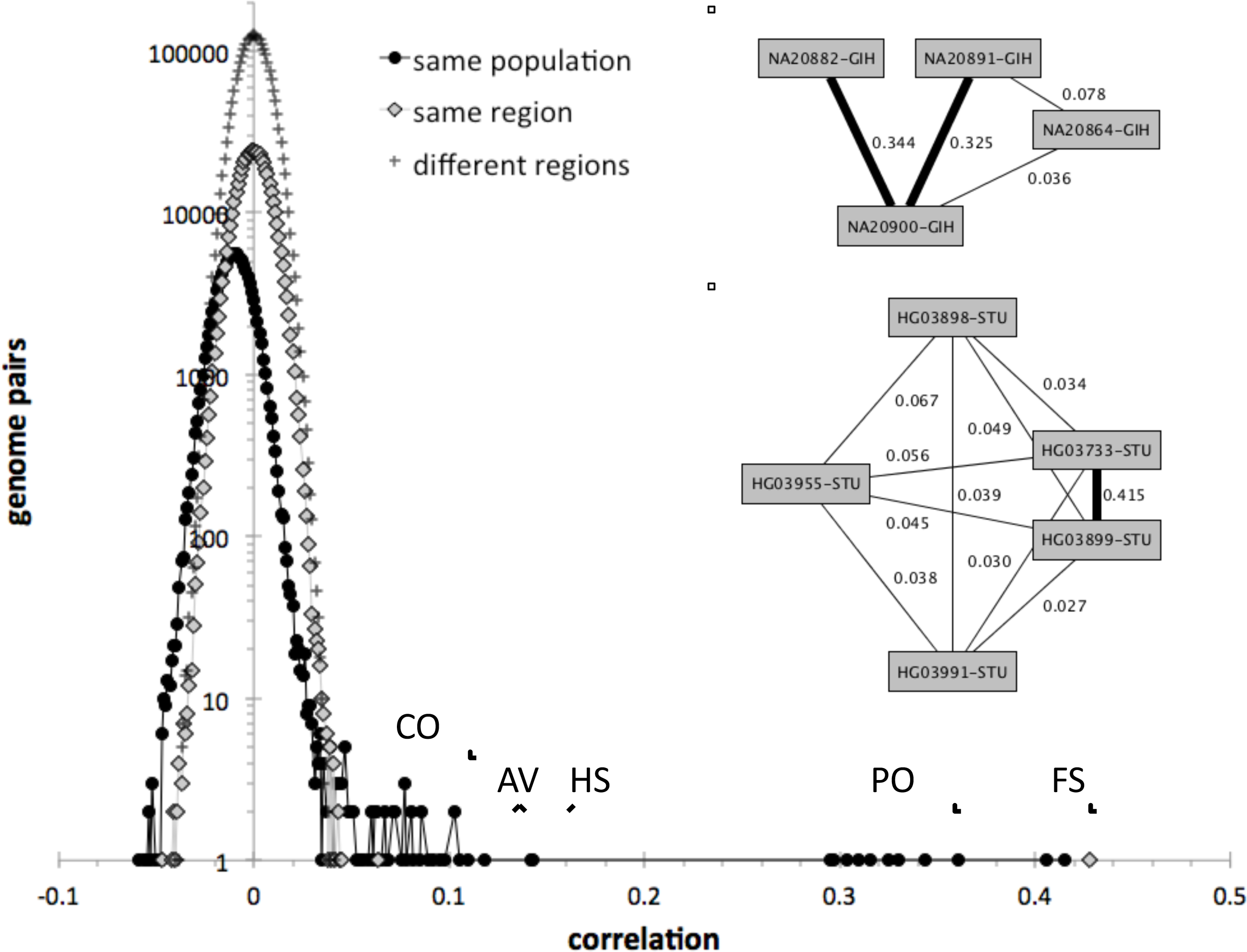
Identification of close relationships in the 1000 Genomes Project. Distribution of correlations of population-adjusted fingerprints for all pairwise comparisons, stratified by whether the two individuals being compared are in the same population, in different populations from the same continental region, or from different continental regions. Close relationships are clearly identified as outliers. FS: full siblings. PO: parent/offspring. HS: half siblings. AV: avuncular. CO: cousins. Upper inset: example of a Gujarati Indian (GIH) family trio with a cousin of one of the parents, recognized through fingerprint comparisons. Thick edges represent PO relationships. Lower inset: example of a Sri Lankan Tamil (STU) pair of siblings and three cousins, identified through fingerprint comparisons. The thick edge represents the FS relationship.

We observed 143 pairs of individuals from the same population with correlations higher than expected (at 5% false discovery rate) and two outlier pairs linking individuals of the ITU and STU populations (Supp. Table 1), consistent with previous reports [11]. The highly correlated pairs correspond to a variety of degrees of relationship, from full siblings to cousins. In several cases they form networks of related individuals, e.g., a Gujarati Indian trio with a cousin of one of the parents (Figure 6, upper inset) and a Sri Lankan Tamil pair of full siblings and three cousins (Figure 6, lower inset).

## Discussion

We have presented a novel algorithm for computing ‘fingerprints’ of individual genomes, and various examples of their potential applications. Genome fingerprints are reduced representations that retain information about distances (between genomes) but not functional features, enabling ultrafast comparison of genomes. Importantly, the fingerprints need to be computed just once per genome, not once per comparison. They can also be computed directly from any representation of the genome, regardless of technology, reference version, formatting and encoding choices, and even significant levels of missing data. These features provide for the first time the ability to rapidly compute the level of similarity between genomes regardless of such representational differences.

We described an implementation of the method for whole-genome data and various aspects of comparing whole-genome fingerprints, pairwise and in large sets. Fingerprints can similarly be computed for exome data. The fingerprint computed from an individual’s exome would not be directly comparable to the fingerprint derived from the same individual’s whole genome; it is though possible and very efficient to approximate exome data from whole-genome data, by simple subsetting using bedtools [12].

Even our unoptimized software for pairwise comparison of genomes via fingerprints takes less than a millisecond; using a compiled language or specialized hardware acceleration could potentially significantly improve on this simple implementation. These speeds are already orders of magnitude faster than afforded by current methods, and enable orders of magnitude more comparisons to be routinely performed. As expected from the highly lossy data reduction involved in generating genome fingerprints, their comparison for various applications yields results that are somewhat inferior to direct computation on the entire dataset of all observed variants. The fast approximation afforded by genome fingerprints will be good enough for some applications, and will prioritize more detailed computations where precise results are required. Widespread adoption of this methodology could revolutionize the field of comparative genomics by enabling comparisons on a scale not now attempted due to the time and effort required.

We have shown that regardless of the technology, genome freeze, or encoding used for specific personal genomes, the corresponding fingerprints enable rapid testing of whether two genome representations are derived from the same person or from closely related individuals, rapid identification of the closest among a specified set of candidate populations, and acceleration of population structure studies. Genome fingerprints are also lightweight and restricted to distance information, and therefore suitable for databasing on a broader scale than the personal genomes themselves. Such databases would enable rapid testing of whether a new personal genome has already been observed, rapidly ensuring that a data set includes only unrelated individuals, and rapid testing for the presence of shared genomes in two or more studies, all of which are common steps in the planning and construction of cohorts of genomes for more sensitive, functional analysis. Distances between fingerprints also provide an unbiased scale for selecting pairs of individuals at comparable distances, for example to select individuals from a control set matched to a set of cases at a consistent evolutionary distance.

Genome fingerprints are highly resilient to a variety of important technical issues, including different reference versions, variant normalization and post-processing, different sequencing technologies, and different variant sampling efficiencies (missing data). We observed that significant post-processing of genome data in a large cohort, as done by the 1000 Genomes Project, can lead to reduced similarities with the unprocessed versions of the same genomes. Several processing steps may contribute to this effect: exclusion of problematic ‘blacklisted’ regions, imputation of missing values, changes in genotypes determined through ‘joint calling’, and the increased chance of observing multi-allelic variants - which are excluded from our analysis.

We evaluated the value of the most compact form of fingerprint, binary fingerprints. Despite their simplicity and compactness, binary fingerprints are typically sufficient to establish whether two genome representations are derived from the same individual, and even to distinguish between closely related and unrelated individuals; however, they do not retain enough information to make these determinations when comparing mixtures of unprocessed and highly post-processed versions of genomes.

Our fingerprinting algorithm is a fully deterministic form of locality-sensitive hashing [10], a class of algorithms which has been successfully applied to large-scale sequence comparisons [13], protein classification [14], metagenome clustering [15] and the detection of gene-gene interactions [16]. Locality sensitive hashing methods based on indexing substrings have been used to compare genomes, typically of different species, and other sequence data types. By encoding single-nucleotide differences from a reference sequence, our method focuses on representing distances among individuals of the same species, for whom the base (reference) sequence is identical and need not be indexed as k-mers. Accordingly, the main application of our method is to detect relationships at various levels - from identity to population structure.

We chose to focus on the most common and well-defined type of genome variation - SNVs. Other types of variation, including short insertions and deletions (indels), larger structural variants, and copy-number variants) are significantly less common than SNVs, and are frequently either represented differently in different genomes, not detected, or not reported: we therefore chose to exclude them for the sake of simplicity and consistency in analysis. Our results demonstrate that SNV-based fingerprints contain enough information to support many genome comparison tasks; the other types of variation are not necessary to estimate evolutionary distances within a species, and could be detrimental for the above reasons.

For each SNV, we only consider the nature of the reference allele and the alternate allele as observed in the individual. Our current implementation considers only positions in which an alternate allele is observed, regardless of zygosity: heterozygous SNVs and homozygous-alternate SNVs are treated equally. We evaluated also alternative versions of the method in which SNVs with different zygosity are weighted differently. Some of these alternative versions will be useful in particular contexts, e.g., for fingerprinting genomes in a fully reference-free fashion by considering only heterozygous SNVs. For each pair of consecutive SNVs, we consider the distance separating them along the reference genome. This is unaffected by any intervening indels or other complex variants in the individual.

We also chose to include only SNVs observed in autosomes. Inclusion of variants in the sex chromosomes leads to distorted similarity values. For example, due to shared variants on the X chromosome, a female child would appear to be more closely related to her mother than to her father, or to be as closely related to her grandfather as to her grandfather’s sister.

Most importantly, our algorithm successfully distributes genome information onto a matrix representation that preserves distances without preserving individual, functionally interpretable variants. While some algorithmic aspects of our method may seem somewhat arbitrary, it is simple and efficient to compute. Prior applications of locality sensitive hashing to genomics have demonstrated the utility of quite arbitrary-seeming algorithms. For example, the kernel of a locality sensitive hashing approach to genome-scale assembly of single-molecule sequencing data uses the XORShift pseudo-random number generator to transform k-mer hashes into comparable ‘sketches’ [14]. The lack of relationship between the mathematical transformation used in our algorithm and evolutionary processes is therefore an advantage that enables genetic distance information to be separated from genetic interpretability, rather than a disadvantage.

The focus on local patterns of SNVs offers an additional advantage of our method. To date, the reference genome has been represented as a collection of linear sequences (chromosomes) with absolute coordinates. One disadvantage of this method is that the absolute coordinates change from version to version; another is that the linear representation is not flexible enough to represent the rich diversity in structural variation observed within our species. The genomics community is now developing a new, graph-based reference genome format, which will further devalue the use of global sequence coordinates as the method for matching sequence variants. We expect that genome fingerprints can be computed from graph-based genome representations, which will be highly comparable to fingerprints we currently compute from linear-reference representations.

Public sharing of genome data has been limited by multiple personal privacy and confidentiality considerations. A central risk is the possibility of identifying genetic predispositions to certain diseases or other traits that could affect the individual’s ability to obtain or maintain employment, insurance or financial services, or may carry social stigma, or could lead to other negative effects. Quantifying this risk is difficult since the ability to interpret genomic variants will expand over time with additional research. Our method enables sharing enough information about a genome to enable the comparison tasks, without concomitantly revealing the information needed for predicting phenotype. As such, our method *decouples genome comparison from interpretation*. This property has important implications for privacy-preserving genome analytics. Identifying genomes harboring a specific variant can be of interest both in the presence of associated phenotype, as facilitated through the Matchmaker Exchange [18], or in its absence, to ascertain novelty and frequency ([19] and the Beacon Network, http://ga4gh.org/#/beacon). Genome fingerprints support the complementary task of matchmaking via identification of closely related individuals, without exposing variant information, in similarity with the UNICORN method [6] but with a much simpler algorithm that doesn’t require extensive prior modeling of variant frequencies, nor samples to be expressed relative to the same reference. Genome fingerprints can furthermore be used to compare genomes from populations not previously studied.

Finally, as growing contingents of private individuals get access to their own genetic data, there is increasing public interest in efficient and private analysis of personal genomic variation, not just for interpreting variants and their combinations, but also for identifying related individuals. As such, our method has strong potential for empowering citizen science.

### Availability

Documentation, code, sample datasets and more are available at: http://db.systemsbiology.net/gestalt/genome_fingerprints

## Acknowledgements

We wish to thank Chris Witwer, Ben Heavner, Irit Rubin, Jeremy Protas and Terry Farrah for helpful discussions. This work was supported by NIH grant U54 EB020406.

## Author contributions

GG, LH and MR designed the study. GG, DM and MR performed analyses. All authors contributed to writing the manuscript and approved its final version.

